# Short-term instantaneous prophylaxis and efficient treatment against SARS-CoV-2 in hACE2 mice conferred by an intranasal nanobody (Nb22)

**DOI:** 10.1101/2021.09.06.459055

**Authors:** Xilin Wu, Yaxing Wang, Lin Cheng, Fengfeng Ni, Linjing Zhu, Sen Ma, Bilian Huang, Mengmeng Ji, Huimin Hu, Yuncheng Li, Shijie Xu, Haixia Shi, Doudou Zhang, Linshuo Liu, Waqas Nawaz, Qinxue Hu, Sheng Ye, Yalan Liu, Zhiwei Wu

**Author notes:** These authors contributed equally to this work (Xilin Wu, Yaxing Wang, Lin Cheng, Fengfeng Ni, Linjing Zhu). Corresponding author: Z. Wu, S. Ye, and Y. Liu, Mailing address: 22 Hankou Road, Nanjing, Jiangsu 210093. China.

## Abstract

Current COVID-19 vaccines need to take at least one month to complete inoculation and then become effective. Around 51% global population are still not fully vaccinated. Instantaneous protection is an unmet need among those who are not fully vaccinated. In addition, breakthrough infections caused by SARS-CoV-2 are widely reported. All these highlight the unmet needing for short-term instantaneous prophylaxis (STIP) in the communities where SARS-CoV-2 is circulating. Previously, we reported nanobodies isolated from an alpaca immunized with the spike protein, exhibiting ultrahigh potency against SARS-CoV-2 and its variants. Herein, we found that Nb22, among our previously reported nanobodies, exhibited ultrapotent neutralization against Delta variant with an IC_50_ value of 0.41 ng/ml (5.13 pM). Furthermore, the crystal structural analysis revealed that the binding of Nb22 to WH01 and Delta RBDs both effectively blocked the binding of RBD to hACE2. Additionally, intranasal Nb22 exhibited protection against SARS-CoV-2 Delta variant in the post-exposure prophylaxis (PEP) and pre-exposure prophylaxis (PrEP). Of note, intranasal Nb22 also demonstrated high efficacy against SARS-CoV-2 Delta variant in STIP for seven days administered by single dose and exhibited long-lasting retention in the respiratory system for at least one month administered by four doses, providing a means of instantaneous short-term prophylaxis against SARS-CoV-2. Thus, ultrahigh potency, long-lasting retention in the respiratory system as well as stability at room-temperature make the intranasal or inhaled Nb22 to be a potential therapeutic or STIP agent against SARS-CoV-2.

**Brief summary:** Nb22 exhibits ultrahigh potency against Delta variant in vitro and is exploited by crystal structural analysis; furthermore, animal study demonstrates high effectiveness in the treatment and short-term instantaneous prophylaxis in hACE2 mice via intranasal administration.

**Highlights:**

1. Nb22 exhibits ultrapotent neutralization against Delta variant with an IC_50_ value of 0.41 ng/ml (5.13 pM).
2. Structural analysis elucidates the ultrapotent neutralization of Nb22 against Delta variant.
3. Nb22 demonstrates complete protection in the treatment of Delta variant infection in hACE2 transgenic mice.
4. We complete the proof of concept of STIP against SARS-CoV-2 using intranasal Nb22 with ultrahigh potency and long-lasting retention in respiratory system.

**Graphic Abstract:** 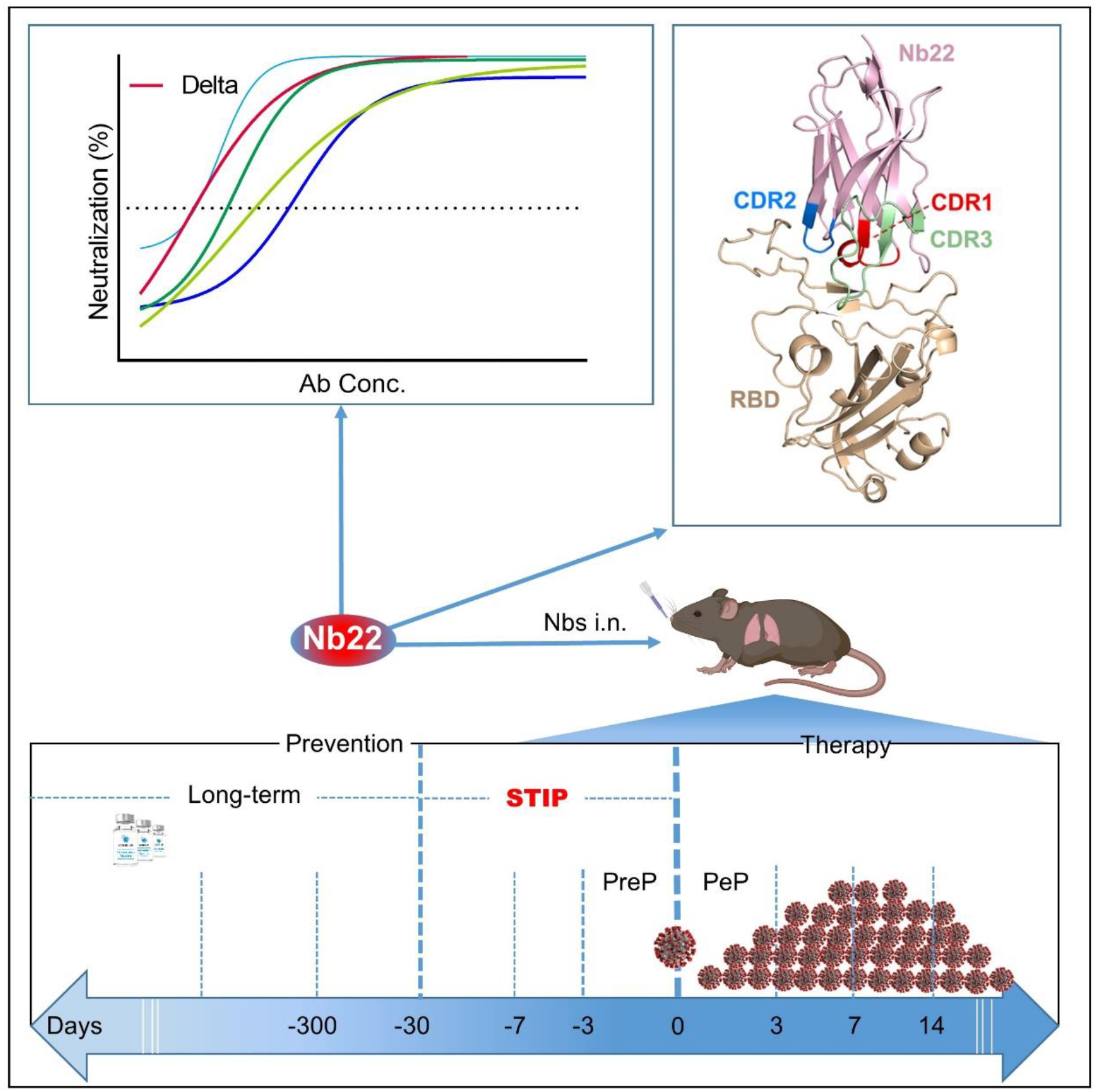

## Introduction

SARS-CoV-2 has given rise to the COVID-19 pandemic ^1^, resulting in massive disruption of social and economic activities. Global vaccination has provided protection against the catastrophic outcome of the pandemic. However, a number of individuals are either not fully vaccinated or cannot mount adequate responses to the vaccine. Additionally, current COVID-19 vaccines require multiple doses to achieve full effectiveness and the immunity wanes within a matter of months, which increases the risk of infection and demands the use of agents for providing instantaneous protection at vulnerable times. Several antibodies were approved for emergency use within 7 days of high-risk exposure in the Post-exposure prophylaxis (PEP) against SARS-CoV-2 infection ^2,3^. However, there is no licensed agent in preventing infection before exposure to SARS-CoV-2 (i.e., as pre-exposure prophylaxis, PrEP). A few PrEP studies in animal model demonstrated that antibodies exhibited accelerated clearance of SARS-CoV-2 when administered 1-3 days prior to infection^2,4–6^. The efficacy was not fully explored when antibodies were administered more than three days prior to SARS-CoV-2 infection. To the best of our knowledge, there is no effective intervention to prevent SARS-CoV-2 infection in advance of one week or longer. Therefore, there is a research gap on short-term instantaneous prophylaxis (STIP) that prevention can take effective immediately following antibody infusion and last for one week or longer. As such, STIP is an unmet need for the prevention against SARS-CoV-2.

The Delta variant, also known as B.1.617.2, was first identified in India in December 2020 and has become predominant in many countries, characterized by the spike protein mutations T19R, L452R, T478K, D614G, P681R, D950N and a double deletion at 157-158 ^7–11^. It has been designated as a Variant of Concern (VOC) and is believed to be 60% more transmissible than Alpha variant ^12^. Delta variant poses a challenge to the available COVID-19 vaccines, such as the protective effectiveness of AstraZeneca and Pfizer vaccines against Delta variant was reduced to 60% and 88%, respectively ^11,12^. More recently, a newly emerged variant, Omicron, has spread rapidly in parts of the world and drawn attention for its potential impact on the global public health; however, in most of the world including China, Delta variant remains the dominant virus and the focus of the containment efforts. Recent research indicated that the Delta variant partially but significantly resisted neutralization by mAbs including Bamlanivimab, SARS-CoV-2 convalescent sera and vaccine-elicited antibodies ^13,14^. While B1-182.1 and A23-58.1, recently isolated from convalescent donors, exhibited ultrapotent neutralization against Delta variant with IC_50_ values of 1.0 and 1.6 ng/ml, respectively ^15^.

To date, a growing number of nanobodies, single-domain fragments of camelid heavy-chain antibodies or VHH, were reported for the prophylaxis or treatment of SARS-CoV-2 infection. However, nanobodies with potent neutralization against Delta variant were rarely reported ^6,16–22^. As SARS-CoV-2 is transmitted through and replicates in respiratory tract and lungs, and does not transverse in blood^1,23^, we believe that, nanobodies’s exceptional resistance to extreme pH and high temperature^24^, makes them ideal candidates to be administered via intranasal or oral rout, directly to the site of viral infection. Previously, we identified three ultrapotent nanobodies against the initial strain of SARS-CoV-2, Wuhan-Hu-01 (WH01); accordingly, one of the intranasally delivered nanobody was shown to protect hACE2 mice infected by WH01 strain^25^.

Here, we compared the neutralizing potency of the aforementioned nanobodies against various circulating SARS-CoV-2 variants. Nb22-Fc was identified to exhibit increased neutralization potency against Delta variant compared to WH01 strain, to which the antibody was originally raised. The binding characterization and crystal structural analysis were conducted to further elucidate the potential mechanism. Furthermore, therapeutic studies demonstrate that intranasal Nb22 exhibited complete protection against SARS-CoV-2 Delta variant in hACE2 mice. Additionally, we comprehensively evaluated the prophylactic efficacy of Nb22 when intranasally administered at 1, 3, 5, 7 days prior to SARS-CoV-2 infection. Notably, single dose of intranasal Nb22 still exhibited protective against hACE2 mice even when administered 7 days prior to infection of Delta variant. Moreover, four doses of intranasal Nb22 could maintain long-lasting retention in respiratory system for more than one month. All these indicate that intranasal Nb22 could be applied not only in the PrEP and PEP but also in the STIP, filling the gap between the long-term lagging prevention and PreP.

## Results

### Potent neutralization of Delta variant by nanobodies

We previously reported the discovery and characterization of three potent neutralizing nanobodies against WH01 strain and its variants with IC_50_ values of ~1 ng/ml. These three nanobodies (Nb15-Fc, Nb22-Fc and Nb31-Fc) were identified to bind to RBD ^25^. Neutralization experiments were further conducted to measure their activity against the circulating variants including variants of concern (VOC) comprising Alpha (B.1.1.7 with N501Y), Beta (B.1.351 with E484K and N501Y), Delta (B.1.617.2 with L452R and T478K) and Gamma ( P.1 with K417T, E484K and N501Y), as well as variants of interest (VOI) comprising Eta (B.1.525 with E484K), Iota (B.1.526 with E484K), Epsilon (B.1.429 with L452R), and Kappa (B.1.617.1 with L452R and E484Q) ^7–10^. Nb15-Fc exhibited increased potency against Alpha variant, but decreased potency against Delta variant or Epsilon as compared with WH01, the RBD used to select the nanobodies. Nb31-Fc exhibited reduced potency against Alpha, Delta and Epsilon variants relative to WH01 or D614G variant (Fig. 1A-G). Interestingly, Nb22-Fc exhibited about 2.5-fold increased neutralizing potency against Delta variant with an IC_50_ value of 0.41 ng/ml (5.13 pM) compared to WH01 with an IC_50_ of 1.01 ng/ml (12.63 pM). Notably, Nb22-Fc also exhibited around 8.4-fold increased neutralization potency against Delta variant relative to variant Alpha with an IC_50_ of 3.45 ng/ml (43.13 pM) (Fig. 1A-G). Impressively, Nb22-Fc also exhibited outstanding neutralizing curve against authentic Delta variant compared to Nb15-Fc and Nb31-Fc (Fig. 1F-G). All three nanobodies failed to neutralize variants containing E484K/Q mutation, suggesting that E484K/Q mutation in RBD could lead to the resistance to all three nanobodies. Altogether, Nb15-Fc presented the most potent neutralization against variant Alpha with an IC_50_ of 0.18 ng/ml, and Nb15-Fc and Nb31-Fc still retained potent neutralization of variants containing L452R and T478K mutations in RBD (Fig. 1E-F), though with reduced potency like most other anti-RBD antibodies ^7,12^. Of note, Nb22-Fc exhibited the most potent neutralization against pseudotyped or authentic Delta variant virus among three nanobodies (Fig. 1A-G).

**Figure 1.**
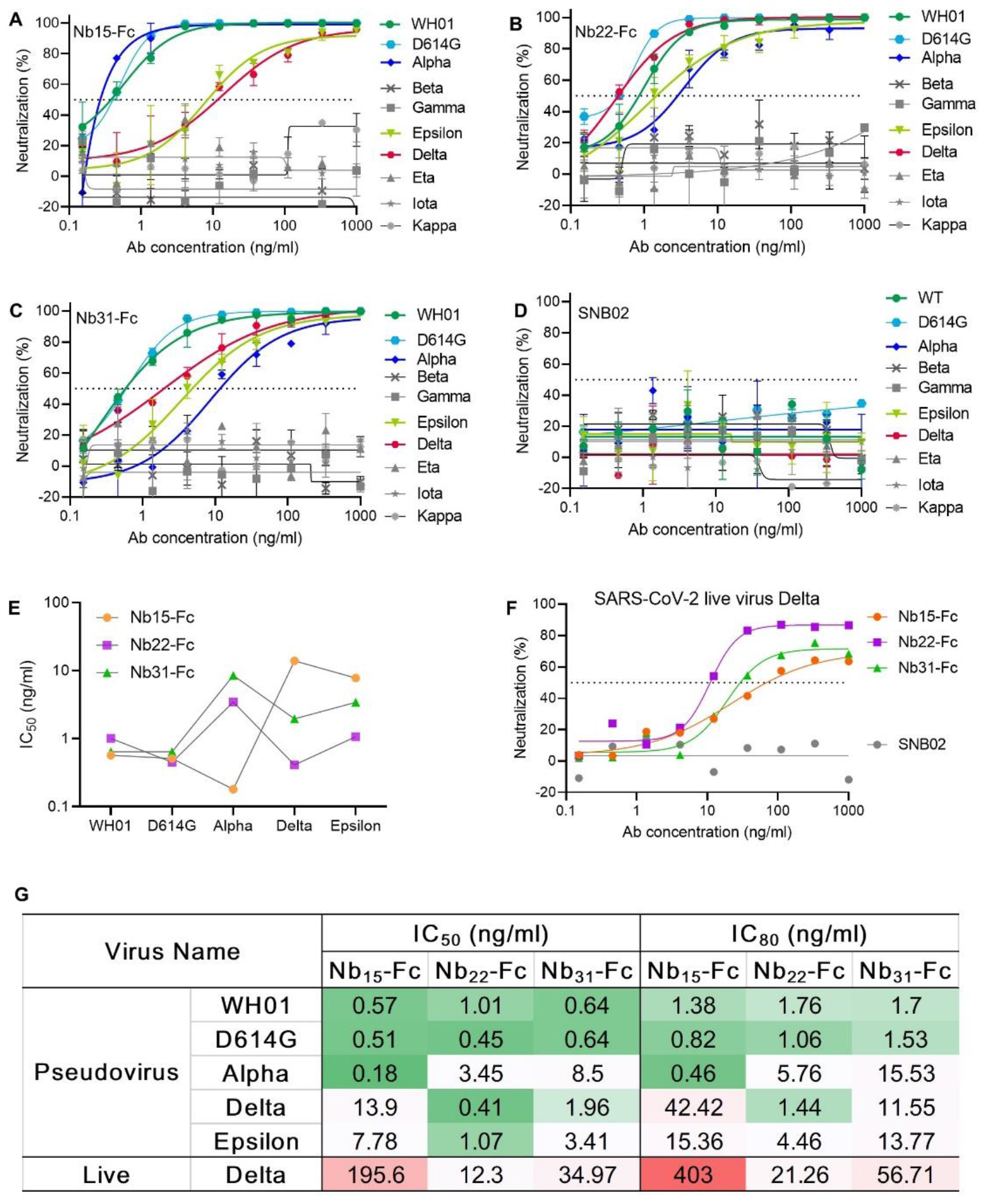
Characterizing nanobodies neutralizing circulating variants of SARS-CoV-2. The neutralization curve of Nb15-Fc (**A**), Nb22-Fc (**B**), Nb31-Fc(**C**) and SNB02 (**D**) inhibiting SARS-CoV-2 pseudovirus of circulating variants. Nb-Fcs and SNB02 were all constructed as the format of VHH fused with human Fc1. SNB02 was taken as an antibody control specific for SFTS virus. (**E)**The summary curve of IC_50_ of Nb-Fcs exhibiting potent neutralization against SARS-CoV-2 variants. (**F**) The neutralization potency of Nb-Fcs was evaluated based on authentic SARS-CoV-2 Delta variant plaque reduction neutralization test. (**G**) The summary table of IC_50_ and IC80 of Nb-Fcs in A-C and F, displaying potent neutralization. Data are represented as mean ± SD. All experiments were repeated at least twice.

### Characterization of Nb22-Fc binding to RBD

To explore antibody binding characteristics to the RBD with respect to their neutralization of Delta variant, the interactions of three nanobodies with variant RBDs were analyzed using biolayer interferometry (BLI). Nb15-Fc, Nb22-Fc and Nb31-Fc showed high affinity interactions with RBD of Delta variant at 1.86 nM, 0.31 nM and 0.31 nM, respectively (Fig. 2A-D). However, the ultrahigh affinity of Nb22-Fc and Nb31-Fc to the RBD of Delta variant did not fully reflect the neutralization potency as Nb22-Fc neutralized Delta variant with markedly more potency than that of Nb31-Fc, suggesting that affinity is not the only factor dictating the neutralization activity. Furthermore, Nb22-Fc exhibited increased affinity with Delta variant RBD relative to other variant RBDs (Fig. 2A-D), which is in line with the increased potency conferred by Nb22-Fc against Delta variant as compared with other variants.

**Figure 2.**
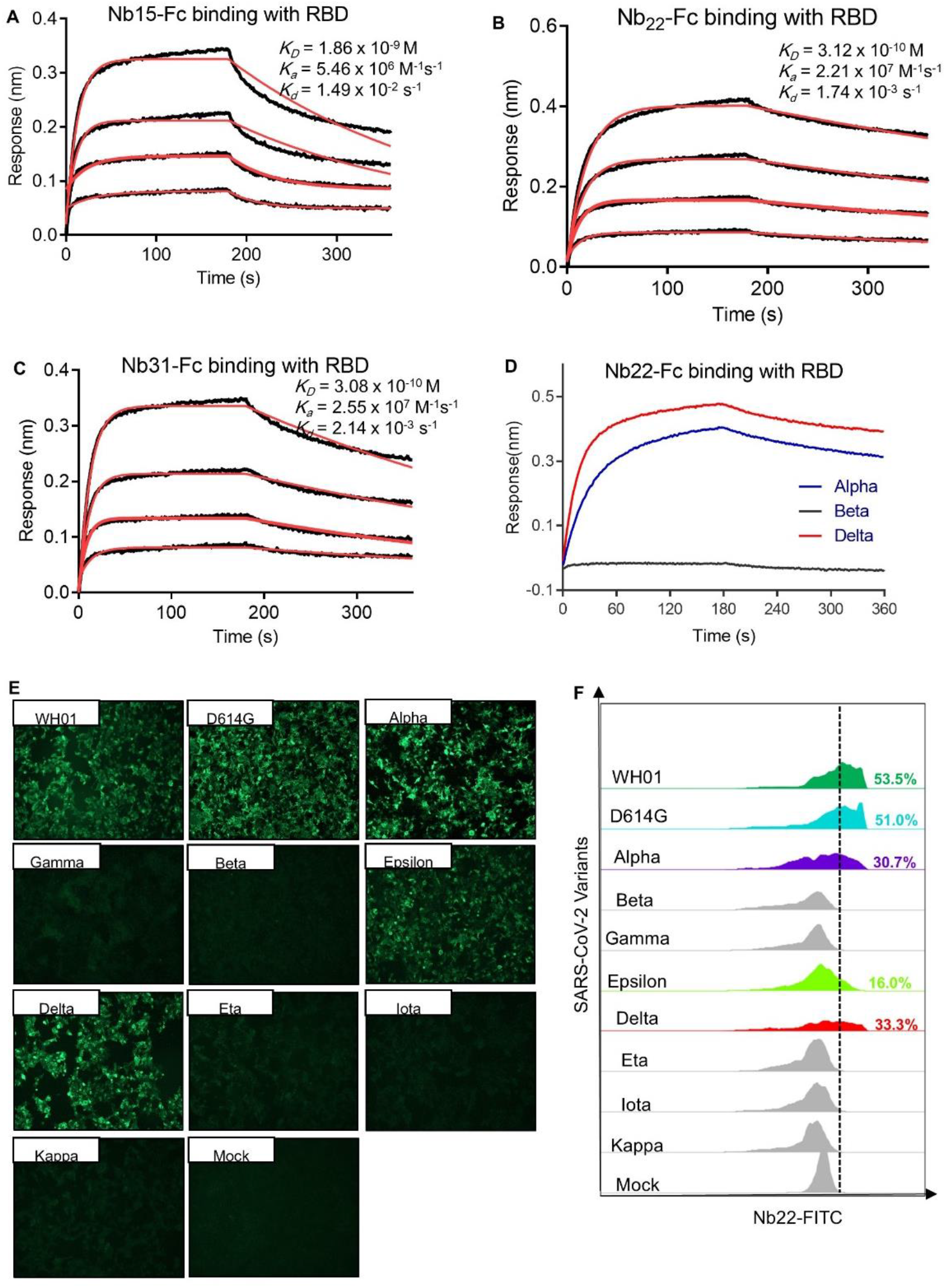
Characterizing the binding of Nbs. Kinetic binding curve of Nb15-Fc (**A**), Nb22-Fc (**B**) and Nb31-Fc (**C**) at the concentration 33.3 nM,11.1nM, 3.7nM and 1.2 nM with RBD of Delta variant, respectively, detected by BLI. Binding curves are colored black, and fit of the data to a 1:1 binding model is colored red. (**D**) Representative binding curve of various RBD as indicated to Nb22-Fc tested by BLI. Nb22-Fc binding with RBD from representative SARS-CoV-2 variants detected by immunofluorescence assay (**E**) and flow cytometric analysis (**F**), respectively. Mock served as a cell control without plasmid transfection. Images were visualized under the ×10 objective. All experiments were repeated at least twice.

Moreover, immunofluorescence analysis revealed that Nb22-Fc specifically interacted with spike protein from WH01, D614G, Alpha, Epsilon and Delta variants on the surface of transfected 293T cells, whereas no binding with the spike protein from other variants containing E484K/Q mutation (Fig. 2E). These results were substantiated by flow cytometric results (Fig. 2F). Overall, these specific binding characteristics are consistent with its specific neutralization properties.

### Structural analysis of RBD-Nb22 complex

Structural analysis of Nb22 interaction with RBD was performed to address the ultrahigh potency of Nb22 against WH01 strain and Delta variant. Initially, we determined the crystal structure of WH01 RBD-Nb22 complex at a resolution of 2.7 Å (Fig. 3A and Table S1). Nb22 adopts a typical β-barreled structure, and contains three variable complementarity-determining regions (CDR) binding to RBD. The buried surface area (BSA) was 800 Å^2^, mainly constituting of hydrogen bonds and hydrophobic interactions. 14 residues constituting epitope of three CDRs were identified using a distance of <4 Å as the cutoff (Fig. 3B). For CDR1, T30 and S33 formed two hydrogen bonds with S494 of RBD, while the hydrophobic interactions included A32 and F34 of Nb22 and Y449, L452, F490 and Q493 of RBD (Fig. 3C and Table S2). N57 of CDR2 interacted with G485 by hydrogen bond, and the hydrophobic interactions were mediated by I56 and Y489 (Fig. 3D). CDR3 is a relatively longer region with only one hydrogen bond (Y119 and G446). The side chain of P104 inserted into the hydrophobic cavity formed by F101, R107, Y453, F456 and Y495 (Fig. 3E). Apart from the five hydrogen bonds in CDR regions, the interface of Nb22 and RBD was stabilized by two additional hydrogen bonds consisting of G1, S75, N450 and E484 (Fig. 3F). Interactions were also facilitated by the hydrophobic network constituted by P2, Q3, V4, G28, G29, R73 and D74 of Nb22 (Fig. 3G and Table S2).

**Figure 3.**
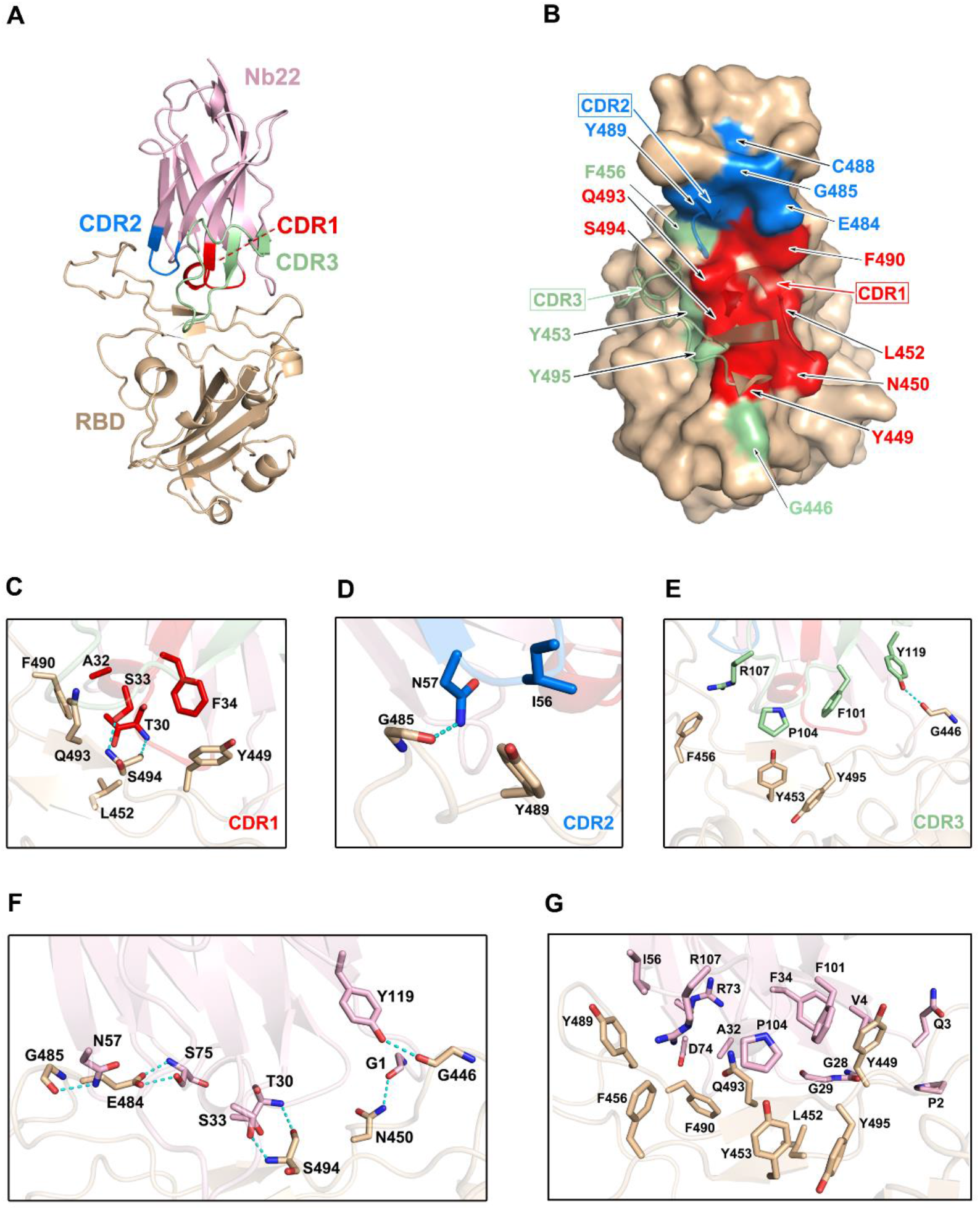
Structural analysis of Nb22 and WH01 RBD complex. (**A**) The overall complex structure of Nb22 and WH01 RBD. The CDR1 (red), CDR2 (blue), CDR3 (green) of Nb22 (pink) and WH01 RBD (orange) are displayed in cartoon representation. (**B**) The epitope of Nb22 shown in surface representation. The CDR regions are colored in red, blue and green, respectively. The interaction between CDR1 (**C**), CDR2 (**D**), CDR3 (**E**) and WH01 RBD. (**F**) The hydrogen bonds of the interface between Nb22 and WH01 RBD. The hydrogen bonds are shown in cyan dash line. (**G**) The hydrophobic network between Nb22 and WH01 RBD. All the residues are shown in sticks.

Superimposition of the structure of WH01 RBD-Nb22 complex and RBD-hACE2 (PDB code: 6MOJ) immediately elucidates the structural basis of neutralization, in which the binding of Nb22 to RBD effectively blocks the binding of RBD to hACE2 during virus infection. Firstly, the binding site of Nb22 on RBD partially overlaps with that of hACE2 (Fig. S1A). Secondly, the loop (V102-Y117) of Nb22 clashes with two α-helices of the N-terminus of hACE2 (Fig. S1B).

To elucidate the increased potency of Nb22 in neutralizing Delta variant, we determined the Delta RBD-Nb22 complex structure at a resolution of 2.9 Å, which revealed that two distinct mutations, T478K and L452R, had differing impact on the binding K478 locates outside the CDR binding regions, and does not disturb the interaction surface (Fig. 4A and Table S1). Therefore, T478K mutation does not affect Nb22 neutralization (Fig. 4A). However, the mutation of hydrophilic leucine to positively charged arginine (R) at position 452 significantly enhances the interactions of RBD with the CDR3 region of Nb22. Two additionally formed hydrogen bonds, T30-R452 and S33-Q493, pull the CDR3 loop of Nb22 closer to R452 region of RBD, as revealed by the superimposition of the structures of WH01 and Delta RBD-Nb22 (Fig. 4B, 4C and Table S2), explaining enhanced neutralization activity of Nb22 against the Delta variant.

**Figure 4.**
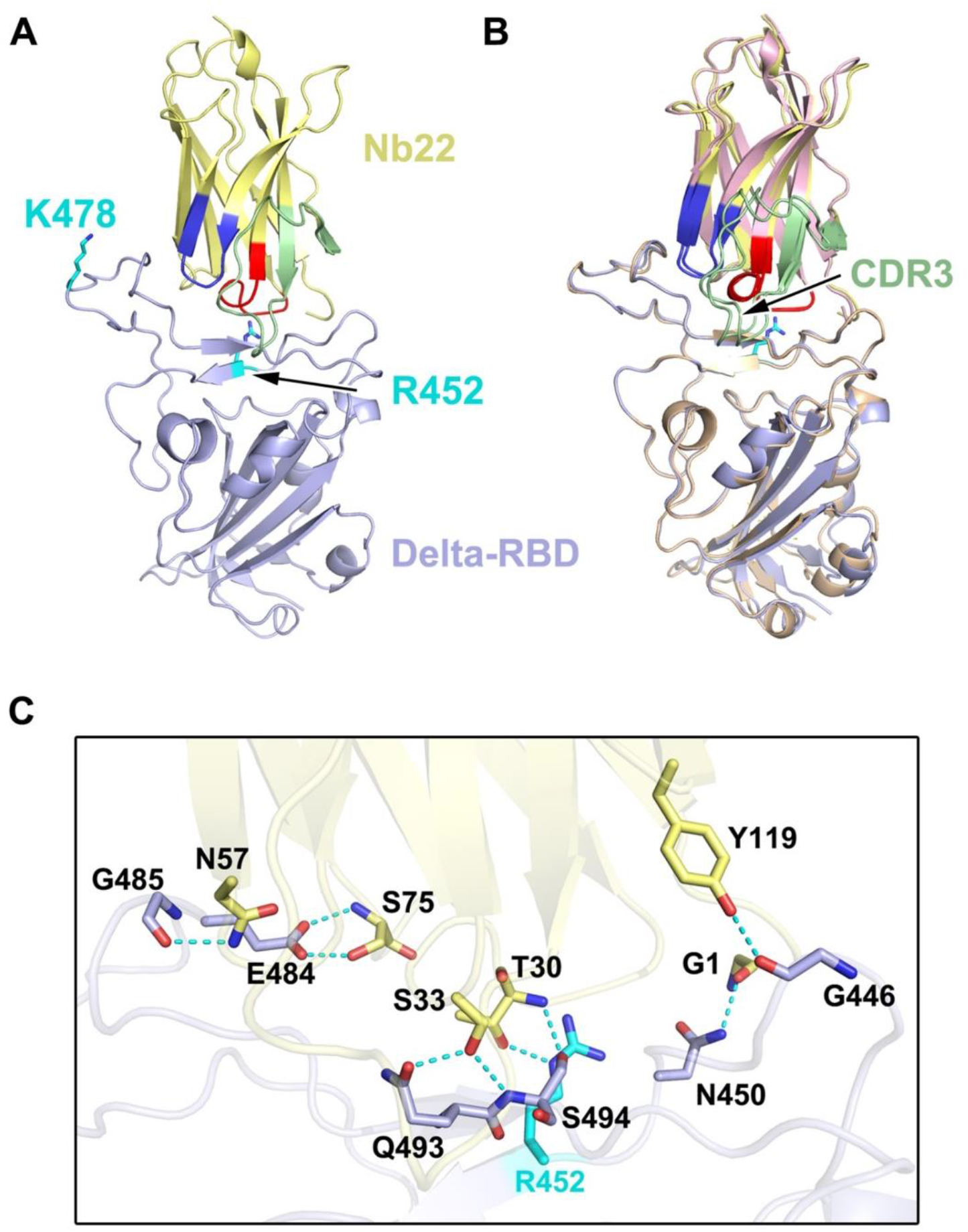
Structural analysis of Nb22 and Delta RBD complex. (**A**) The two mutation sites of Delta RBD. K478 is located outside the CDR binding regions, R452 is on the CDR2 recognized epitope. R452 and K478 are colored in cyan, and the epitope of CDRs is colored identical to Fig. 3. **(B)** The superimposition of WH01 RBD-Nb22 (orange and pink) and Delta RBD-Nb22 (light blue and yellow). **(C)** The hydrogen bonds on the interaction interface of Delta RBD-Nb22. R452 and Q493 form two additional hydrogen bonds with T30 and S33 of Nb22 The residues identified are shown in sticks.

### Nb22 exhibits room-temperature stability in vitro and long-lasting retention in vivo

Nanobodies exhibit various advantages including thermostability. We reported that nanobodies could retain 100% activity even after being incubated at 70 °C for one hour and retain 83% activity after 80 °C treatment for one hour^25^. Further evaluation showed that Nb22 could maintain full activity for at least two months at room temperature and did not lose any activity even undergoing five rounds of freeze-thawing (Fig. 5A and 5B), indicating that Nb22-Fc is highly stable and idea for non-cold chain storage and use at room-temperature.

**Figure 5.**
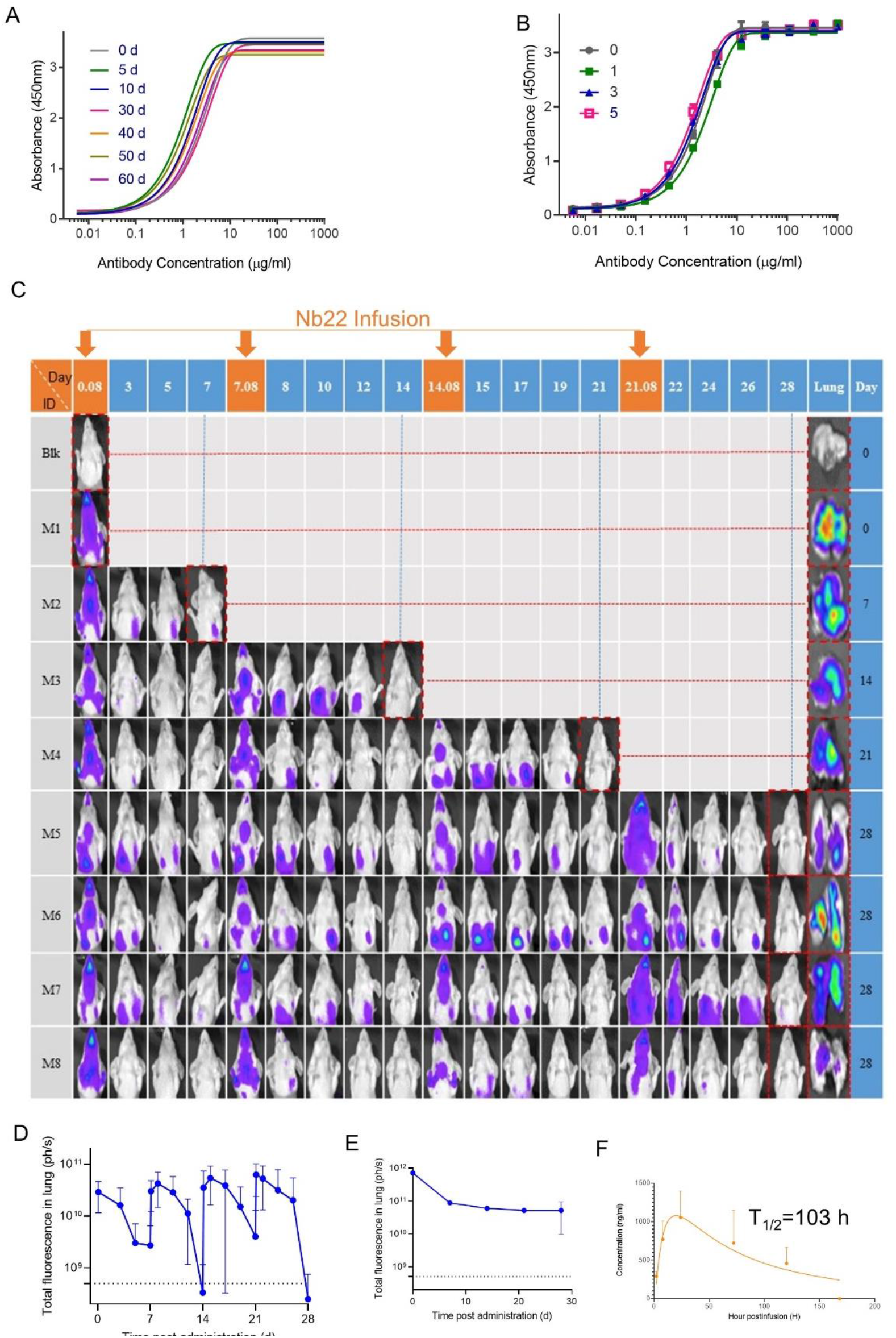
Characterizing Nb22-Fc stability in vitro and pharmaceutics in vivo. (**A**) Binding curve of RBD with Nb22-Fc detected by ELISA after storage at room temperature in the indicated time points including 0 d, 5 d, 10 d, 30 d, 40 d, 50 d and 60 d. (**B**) Binding curve of RBD with Nb22 detected by ELISA after the indicated rounds of freeze-thawing including 0, 1, 3, 5 rounds, respectively. (**C**) Pharmacokinetic of Nb22-Fc labeled with dye YF^®^750 SE via intranasal administration was detected. 200 μg Nb22-YF750 was infused at day 0, 7, 14 and 21 respectively. The optical imaging of mice upper half body was measured by NightOwl LB 983 0.08 d (2h) post Nb22-Fc infusion or at indicated time point labeled at the top of panel. The mice in the red dash line figure were sacrificed at the indicated time point in the left of panel for analysis of the fluorescence intensity of lung. (**D**)The fluorescence intensity of upper half body of mice in (**C**) was summarized. (**E**)The fluorescence intensity of lung in lung column of (C) was summarized. (**F**) Bioavailability and t1/2 of Nb22 in BALB/c mice.Nb22 was intranasally (i.n.) administered into mice (n=3, Female) at 200 μg (average of 10 mg/kg mice. Serum concentrations of the Nbs were determined at various time points by ELISA. T1/2, time of half-life. Data represent mean ± SEM.

To determine Nb22-Fc distribution in vivo, YF^®^750 SE-labeled Nb22 (Nb22-YF750) was administered via intranasal (i.n.) in a mouse model. The fluorescence could be readily detected in respiratory system including nasopharynx, trachea and lung 2h post infusion. The fluorescence was detectable for more than seven days after a single dose of 200 ug (average of 10 mg/kg) Nb22-YF750 administration, which is in agreement with our previous reports. As expected, the fluorescence could be detected for more than one month when four doses of 200 μg (average of 10 mg/kg) Nb22-YF750 were administered every week (Fig. 5C-E and Fig. S2). Nb22-Fc could also be detected in the blood, indicating that Nb22-Fc could also exert its activity in the blood after bypassing the respiratory system (Fig. 5F). All these observations of prolonged retention of Nb22-Fc in the respiratory system suggest the potential application of the antibody for STIP against SARS-CoV-2 infection. Taken together, intranasal Nb22-Fc could be developed as an STIP reagent for its long-lasting retention in the respiratory system and a portable therapeutics thanks to its room-temperature stability.

### Intranasal Nb22 is highly efficacious in the STIP, PreP and PeP in hACE2 transgenic mice challenged by Delta variant

To evaluate the efficacy of Nb22-Fc in vivo, hACE2 transgenic mice were challenged with 1×10^5^ PFU SARS-CoV-2 Delta variant (CRST: 1633.06.IVCAS 6.7593) and conducted as we previously reported^25^. hACE2 mice were divided into nine groups (n=3-6) as shown in Figure 6A and 200 μg (average of 10 mg/kg) Nb22-Fc was administered via i.n. before or after Delta variant challenge to evaluate the antibody’s prophylactic or therapeutic efficacy against Delta variant infection. Viral RNA in the lungs was detected in virus control group named as SARS-CoV-2 group (Group 2). Animals in Control Nb group Group 3 were challenged with Delta variant and received control nanobody treatment one hour post infection. As expected, high copy numbers of viral RNA were also detected in control Nb mice without significant difference compared to SARS-CoV-2 group (Fig. 6B).

**Figure 6.**
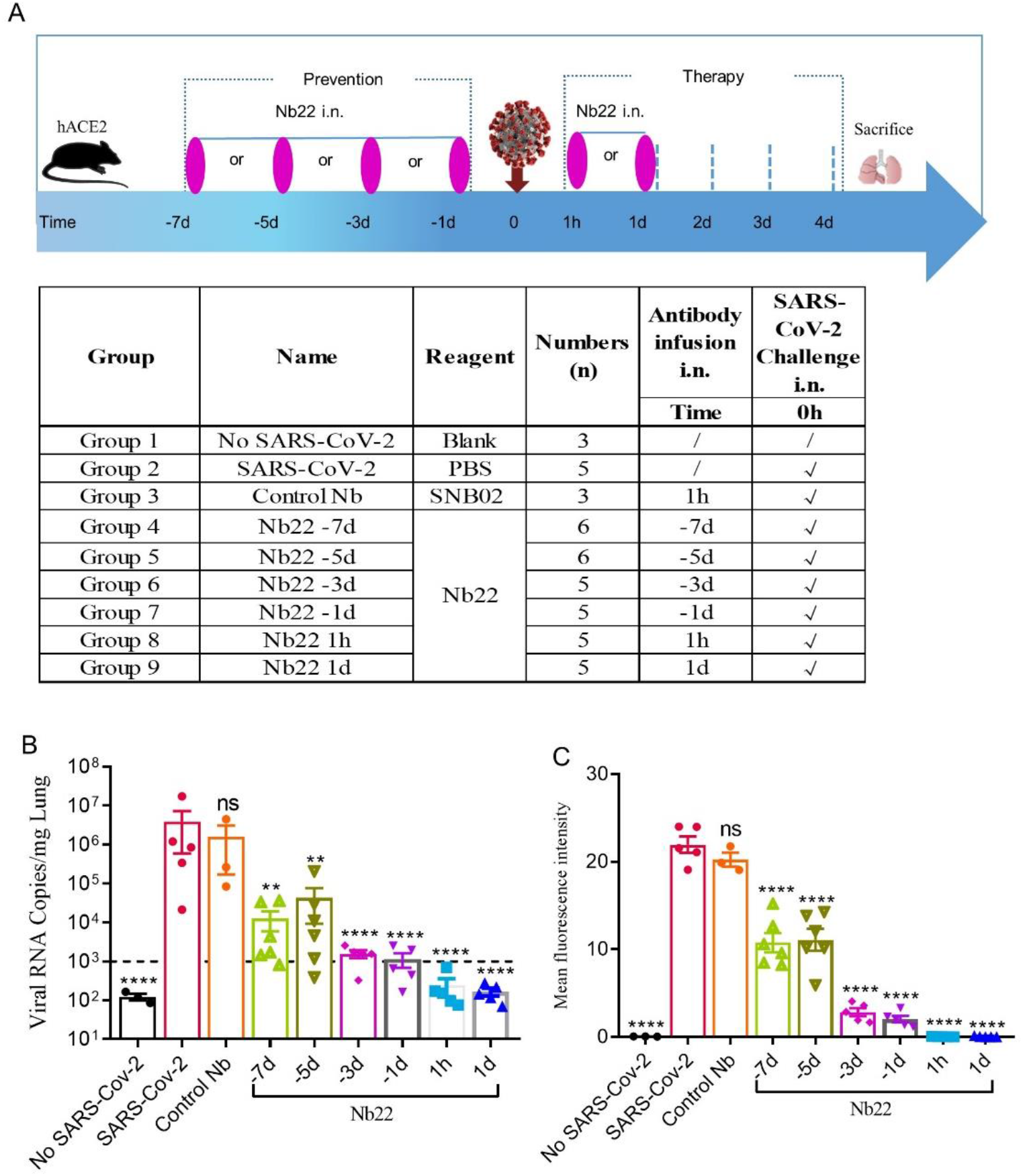
The efficacy of Nb22s evaluated in hACE2 transgenic mice challenged by SARS-CoV-2. (**A**) Experimental schedule of Nb22s in the prevention and treatment of SARS-CoV-2 infection. Bottom, table summary of groups (n = 3–6 mice) with different treatments. (**B**) Viral loads in lungs among 9 groups were measured by qRT-PCR. The name of each group in the x axis was indicated as in the table in (A). Each dot represents one mouse. The limit of detection was 1000 copies/mg referenced to blank control (No-SARS-CoV-2 group). Data are represented as mean ± SEM; Mann - Whitney test was performed to compare treatment group with the SARS-CoV-2 control group. (**C**) Sections of lung were analyzed by immunofluorescence staining using antibodies specific to SARS-CoV-2 NP in red and DAPI for nuclei in blue, respectively. The fluorescence signal intensity of red was taken as a quantitative indicator for viral infection, which was calculated by ImageJ software. ns, no significance; *p < 0.05, **p < 0.01, ***p < 0.001. All experiments of (B) and (C) was repeated twice.

In order to evaluate the prophylactic duration conferred by Nb22, a single dose of Nb22 was administered via i.n. at days 1, 3, 5, and 7, respectively, prior to Delta variant challenge. Viral RNA copies increased over the course of Nb22-Fc administration (Fig. 6B). As expected, viral RNA copies in the aforementioned prophylactic groups were all significantly lower than that in the control Nb group, indicating that even a single dose of Nb22-Fc could provide protection against Delta variant infection in hACE2 transgenic mice for 7 days. Nb22 administered in −7d and −5d before challenge significantly reduced viral load though failed to provide complete protection (Fig. 6B). Notably, Nb22 exhibited significant prevention against SARS-CoV-2 infection in the - 3d Nb22 (Group 6) and −1d Nb22 group (Group 7) (Fig. 6B). All these indicate that Nb22-Fc provides better protection at the earlier time points upon infection.

In the therapeutic group, viral RNA copies in the animals treated with Nb22 at 1 hour or 1 day postinfection were undetectable in 5/5 mice in the groups 8 and 9, suggesting that Nb22 had complete protection of hACE2 mice against Delta variant infection (Fig. 6B). The viral RNA results were also validated by immunofluorescence (IF) staining and HE staining in the lungs (Fig. 6C and 7E). We noted that mice challenged by Delta variant did not show obvious weight loss even in 4 days post infection (Fig. S3). In summary, Nb22 exhibited high efficacy in both the prevention and therapy of hACE2 transgenic mice challenged with Delta variant. Nb22 provided complete protection in PEP (in 1h Nb22 group and 1d Nb22 group) and exhibited high efficacy in PrEP (in - 1d Nb22 and −3d Nb22 group). Impressively, a single dose of Nb22 could maintain effectiveness in the prevention against Delta variant infection for at least seven days (in −7d Nb22 group), indicating the potential application for STIP against SARS-CoV-2.

**Figure 7.**
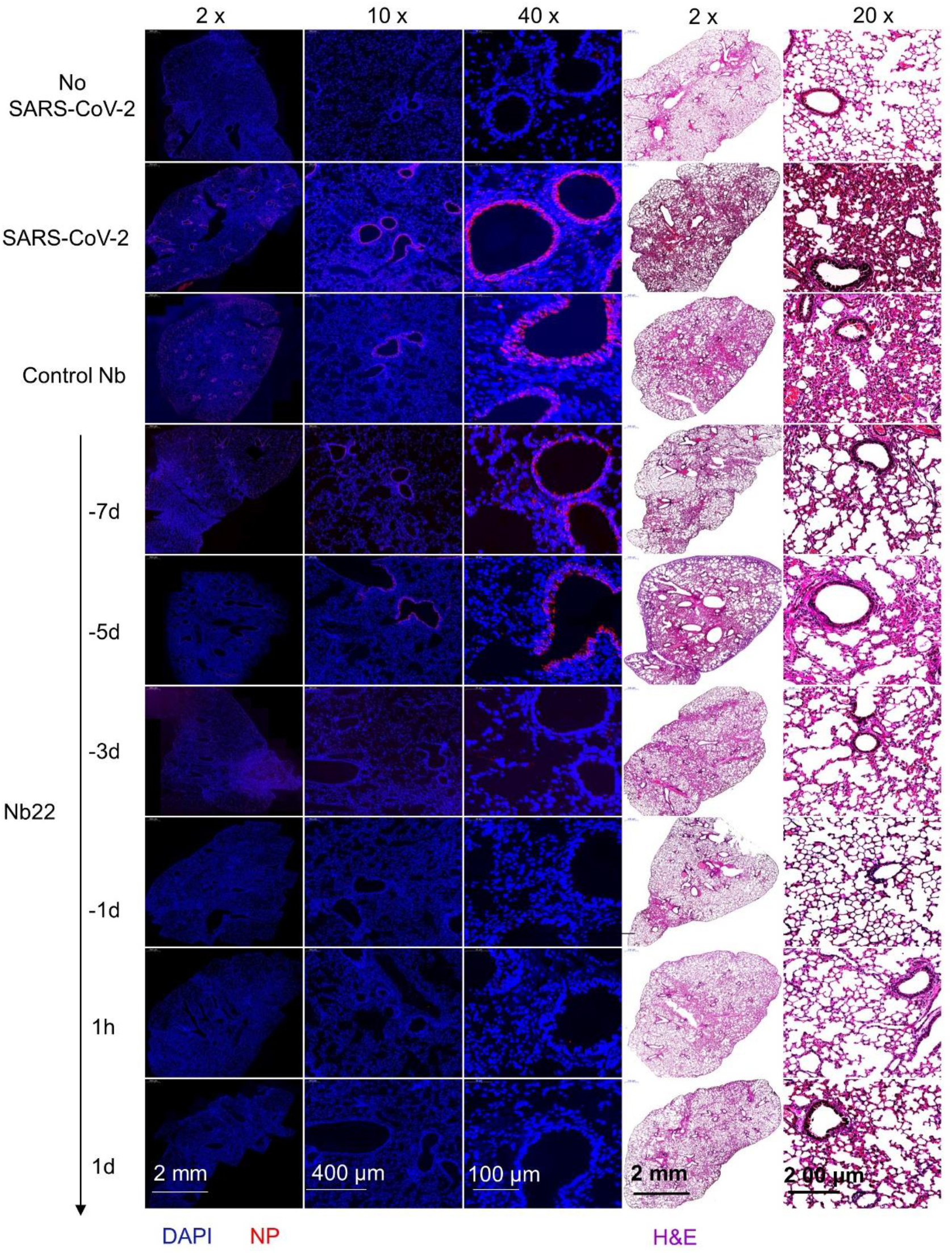
Representative sections of lung from hACE2 mice were analyzed by H&E staining and immunofluorescence staining. Representative lung tissue sections from the mice indicated in figure 6C including No-SARS-CoV-2, SARS-CoV-2, Control Nb, -7d-Nb22, -5d-Nb22, -3d-Nb22, 1h-Nb22 and 1d-Nb22 group were analyzed by immunofluorescence staining using antibodies specific to SARS-CoV-2 NP in red and DAPI for nuclei in blue, respectively. The corresponding representative lung tissue sections were also analyzed by H&E staining. Immunofluorescence and HE Images were visualized under the indicated bar.

## Discussion

To date, a small number of nanobodies with ultrahigh potency against SARS-CoV-2 and its variants have been reported ^15,26,27^, whereas nanobodies with potent neutralization against the currently dominant Delta variant were rarely reported. Our results revealed that three previously reported nanobodies ^25^, retained ultrahigh potency in neutralization against the Delta variant. Among them, Nb22-Fc with an IC_50_ value of 0.41ng/ml (5.13 pM) is outstanding with increased neutralization of Delta variant relative to Alpha variant. The Nb22 binding to RBD provides mechanistic insight into the enhanced neutralization against Delta variant, suggesting that the increased binding affinity enhanced the neutralizing potency against Delta variant relative to Alpha variant (Fig. 1). Given that most anti-RBD, anti-NTD antibodies or convalescent sera or vaccine-elicited antibodies showed reduced neutralization of Delta variant relative to that of Alpha variant ^7,12^, the increased neutralization activity of Nb22-Fc against Delta variant is particularly striking and the structural basis of the phenomenon is of interest for understanding the neutralization mechanisms.

The structural analysis further illustrated the characteristics of Nb22 binding to WH01 and Delta RBD and the mechanisms of viral inhibition. Nb22 binding to RBD effectively blocks the binding of RBD to hACE2 during virus infection. The binding site of Nb22 on RBD overlaps with that of hACE2 (Fig. S1A), and the loop (V102-Y117) of Nb22 clashes with two α-helixes of the N-terminus of hACE2 (Fig. S1B). In addition, crystal structural analysis showed that T478K mutation of Delta variant are located outside 800 Å^2^ BSA of Nb22 interacting with RBD and do not perturb the interaction between Nb22 and the RBD of Delta variant. Of note, the guanidine moiety in the L452R mutation forms an additional hydrogen bond with the hydroxyl group of T30 of Nb22, pulling the CDR3 loop of Nb22 closer, and leading to an extra hydrogen bond between S33 of Nb22 and Q493 of RBD (Fig. 4). Consequently, the BSA extends from 800 Å^2^ to 835 Å^2^, in comparison with that of Nb22-WH01 RBD. All these contribute to the enhanced binding and neutralizing potency of Nb22 against Delta variant.

Compared to conventional antibodies for passive immunization, nanobodies are efficiently produced in prokaryotic expression systems at low cost and possess favorable biophysical properties including high thermostability^28^. We reported that nanobodies could remain 100% activity even incubated at 70 °C for one hour and are amenable to engineering of multimeric nanobody constructs^25^. Such nanobodies exhibit high effectiveness against virus infection via intranasal administration^25^. The results reveal that Nb22 could maintain full activity for more than two months at room temperature and does not lose any activity even after undergoing five rounds of freeze-thawing.

Our results demonstrate that a single dose of intranasal Nb22 could exhibit efficacy in the STIP, PrEP and PEP against SARS-CoV-2 infection in hACE2 mice. Of note, a single dose of intranasal Nb22 could maintain efficacy against SARS-CoV-2 infection for at least one week in hACE2 mice, which would readily serve as STIP. Our antibody distribution results also revealed that Nb22 could retain in respiratory tracts for at least one month when weekly administered via i.n.. As such, we anticipate that Nb22 could provide one month prevention against SARS-CoV-2 infection when administered intranasally every week.

VLong–term lagging prevention against SARS-CoV-2 conferred by approved vaccine usually takes more than one month to be effective and then lasts for months or years^29^. Vaccine efficacy has been shown to wan within months after vaccination ^7,11^. Instantaneous prevention of SARS-CoV-2 is also needed for individuals who do not take vaccines when SARS-CoV-2 is circulating. A few studies in animal model just demonstrated that antibodies exhibited accelerated clearance of SARS-CoV-2 in PrEP when administered 1-3 days prior to infection^2,4–6^. Whereas, to the best of our knowledge, few studies have fully investigated the STIP that prevention could be readily effective immediately following inoculation and last for more than one week to one month for people at high risk of SARS-CoV-2 infection, which can also serve to reduce transmission during the asymptomatic stage of the infection As such, our results demonstrate that intranasal Nb22 with ultrahigh potency and long-lasting retention in the lung could satisfy the need of STIP against SARS-CoV-2.

In summary, structural analysis provides a mechanistic explanation to the enhanced sensitivity of Delta variant and the increased neutralization potency of this antibody. The structural analysis may further guide the rational design of pan-coronavirus vaccines and therapeutics. Nb22 exhibited one of the ultrahigh neutralization potencies among the reported antibodies or nanobodies against Delta variant infection ^7,12,26,30,31^. We presented proof of concept of STIP against SARS-CoV-2 using our Nb22 and suggest STIP as a new prophylactic strategy for long-lasting antibodies to prevent virus infection. Although the newly emerged Omicron variant is spreading globally, the Delta variant remains the dominant variant and likely the deadliest in most of the world; therefore, the ultrahigh potent, and thermal stable Nb22 is an excellent candidate for intranasal or inhalable anti-SARS-Cov-2 agent for both therapy or prophylaxis, especially including STIP.

## Materials and Methods

### Expression of nanobodies

The Fc1 gene (CH2-CH3) of the human monoclonal antibody was fused with the VHH gene of nanobodies (named as Nb-Fc or Nbs) to assist the purification and prolong the half-life of the Nb antibody, following our previously published protocol ^32^. The Nbs were finally cloned into the pcDNA3.4 eukaryotic expression vector (Invitrogen), which were transfected into 293F cells (cat.# R79007, Thermo Scientific) to produce Nb-Fcs. Nb fused with Fc was purified using Protein G (cat.# 20399, Thermo Scientific).

### Enzyme linked immunosorbent assay (ELISA) analysis

Antibody quantification and antibody characterization were tested by ELISA as our previously reported method^33^, with modifications. In brief, the protein was coated to ELISA plates (Corning) at a concentration of 0.5 μg/ml. After washing 2-4 times, 5% non-fat milk in PBS was added and incubated for blocking at 37 °C for 1 h. After washing, 100 μl serially diluted sera or purified antibody was added and incubated at 37 °C for 1 h. Following washing, secondary antibody of goat anti-llama IgG (H+L) with HRP (Novus, cat.# NB7242, 1:10000 dilution) or goat anti-human IgG with HRP was added and incubated at 37 °C for 1 h. Accordingly, 3,3′,5,5′-Tetramethylbenzidine (TMB, Sigma) substrate was added at 37 °C for 10 minutes (min); and 10 μl 0.2 M H_2_SO_4_ was added to stop the reaction. The optical densities at 450 nm (OD450) were measured using the Infinite 200 (Tecan, Ramsey, MN, USA). Antibody quantification in the sera was calculated according to the standard curve generated by purified antibody.

### Neutralization activity of nanobodies against pseudovirus

Pseudovirus neutralization assay was carried out following our previously published protocol^25^, with the follow modifications. Briefly, pseudovirus of SARS-CoV-2 variants was produced by co-transfection of pNL4-3.Luc.R-E-, an HIV-1 NL4-3 luciferase reporter vector that comprises defective Nef, Env and Vpr (HIV AIDS Reagent Program), and pVAX1 (Invitrogen) expression vectors encoding the spike proteins of respective variants into 293T cells (ATCC). Supernatants containing pseudovirus were collected after 48 hours (h), and viral titers were determined by luciferase assay in relative light units (Bright-Glo Luciferase Assay Vector System, Promega Biosciences). Human codon optimized S genes of SARS-CoV-2 variants were synthesized, and the corresponding pseudoviruses were produced following the above protocol. For neutralization assay, SNB02, an Nb-Fc specific against SFTSV ^32^, served as a negative control. Neutralization assays were conducted by incubating pseudovirus with serial dilutions of purified nanobodies at 37 °C for 1 h. HEK293T-ACE2 cells (cat.# 41107ES03, Yeasen Biotech Co., Ltd. China) (approximately 2.5×10 ^4^ per well) were added in duplicate to the virus-antibody mixture. Half-maximal inhibitory concentrations (IC_50_) of the evaluated nanobodies were determined by luciferase activity 48 h following exposure to virus-antibody mixture, and analyzed by GraphPad Prism 8.01 (GraphPad Software Inc.).

### Immunofluorescence and flow cytometric analysis

Immunofluorescence and flow cytometric analysis were conducted following our previously published protocol ^34^, with minor modifications. Briefly, S gene sequences for SARS-CoV-2 spike protein of various SARS-CoV-2 variants were obtained from the GISAID website (https://gisaid.org). S genes were synthesized and constructed as expression plasmids by GenScript. The plasmids were transfected into 293T cells (ATCC) cultured in 12-well plates. Next, 48 hours post transfection, the cells were washed by PBS and fixed with 4% paraformaldehyde for 20 minutes at room temperature. The purified Nb-Fc was used to stain the 293T cells, followed by Alexa Fluor 488 AffiniPure goat Anti-human IgG (H+L) (1:500 dilution) (109-545-003, Jackson ImmunoResearch). For immunofluorescence analysis, the cells on the plate were examined and the images were acquired using an OLYMPUS IX73. For flow cytometric analysis, the cells were resuspended in 500 µl PBSF buffer (PBS+2% FBS) and analyzed using ACEA NovoCyte TM (Agilent Biosciences); non-transfected 293T cells served as a negative control.

### Affinity determination by Bio-Layer Interferometry (BLI)

We measured antibody affinity using a ForteBio OctetRED 96 BLI (Molecular Devices ForteBio LLC, Fremont, CA) with shaking at 1,000 rpm at 25 °C ^25^. To determine the affinity of Nbs with human Fc tag, Nb-Fcs were loaded to anti-human Fc (AHC) biosensors (cat.# 18-5060, Fortebio) in a kinetic buffer (PBS, 0.02% (v/v) Tween-20, pH 7.0) for 200 sec prior to baseline equilibration for 180 sec in a kinetic buffer. Association of SARS-CoV-2 RBD in a three-fold dilution series from 33.3 nM to 1.2 nM was performed prior to dissociation for 180 sec.After each cycle, the biosensors were regenerated through 3 brief pulses of 5 sec each with 100 mM pH 2.7 glycine-HCL followed by a running buffer. The data were baseline subtracted before fitting using a 1:1 binding model and the ForteBio data analysis software. *K*_*D*_, *Ka* and *Kd* values were determined by applying a global fit to all data.

### Expression and purification of WH01 and Delta RBD protein for crystal structural analysis

The WH01 and Delta RBD were expressed using the Bac-to-Bac baculovirus system. The two pAcgp67-RBD (residues 333–530) plasmid with a C-terminal 8×His tag were transfected into Sf9 cells using Cellfectin II Reagent (Invitrogen) to produce the recombinant baculoviruses. After 3 rounds of amplification, Hi5 cells were infected with baculoviruses at an MOI of 4 at a density of 2 × 10^6^ cells/ml. The supernatants of cell culture containing the secreted RBD were harvested at 60 h after infection. The RBD was purified by Ni-NTA resin (GE Healthcare). Nonspecific contaminants were removed by washing the resin with 20 mM Tris-HCl, 150 mM NaCl, pH 7.5, and the target proteins were eluted with elution buffer containing 20 mM Tris-HCl, 150 mM NaCl, 500 mM imidazole, pH 7.5. The eluted proteins were further purified by Superdex 75 (GE Healthcare, USA) and stored in 20 mM Tris-HCl, 150 mM NaCl, pH 7.5.

### Expression and purification of Nb22 for crystal structural analysis

The VHH gene for Nb22 was amplified by PCR and cloned into a pET21a vector with *Bam*H I and *Xho* I restriction sites. The recombinant plasmids were transformed into *Escherichia coli*. BL21 (DE3). The cells were cultured in LB medium and grown to OD_600_ = 0.8 at 37°C. Isopropyl -D-1-thiogalactopyranoside (IPTG) was added to a final concentration of 1.0 mM to induce the protein expression, and the cultures were grown at 16 °C overnight. Cells were harvested by centrifugation at 4,500 rpm for 15 min, re-suspended and homogenized in the lysis buffer containing 20 mM Tris-HCl, 150 mM NaCl, pH 7.5 using ultrasonic. Cell debris was removed by centrifugation at 18,000 rpm for 30 min. The supernatants were added to Ni-NTA resin (GE Healthcare, USA). The nonspecific contaminants were eluted by washing the resin with the lysis buffer containing 10 mM imidazole. The target protein with 6 x His tag, named as Nb22, was subsequently eluted with the lysis buffer containing 500 mM imidazole. Nb22 was eluted and purified by Superdex 75 (GE Healthcare, USA).

### Crystallization, structural determination and data acquisition

The complexes were prepared by mixing WH01 or Delta RBD and Nb22 at a 1:1.2 molar ratio and incubating at 4 °C overnight. The complexes were further purified by Superdex 75 (GE Healthcare, USA) to remove the excess nanobody. The crystals were screened by vapor-diffusion sitting-drop method at 16°C. The crystals appeared and reached their final size within 3 days in a well solution comprising 0.1 M HEPES (pH 7.0), 5% *v*/*v* (+/−)-2-Methyl-2,4-pentanediol (MPD), 10% polyethylene glycol (PEG) 10000 (WH01 RBD-Nb22) and 0.1 M Tris (pH 7.0), 37.5% Jeffamine (Delta RBD-Nb22), respectively.

To collect data, a single crystal was mounted on a nylon loop and was flash-cooled with a nitrogen gas stream at 100 K. Diffraction data of WH01 RBD-Nb22 was collected on BL18U1 at Shanghai Synchrotron Radiation Facility (SSRF) at a wavelength of 0.97915 Å. While, the Delta RBD-Nb22 was collected on BL02U1 at a wavelength of 0.97918 Å. Data were processed and scaled using the HKL3000 package and autoPROC ^35^. The structures were elucidated using the molecular replacement (MR) method in PHASER program ^36^ with the structure of SARS-CoV-2 RBD (PDB code: 7CJF) ^37^ as the initial searching model. The model was built into the modified experimental electron density using COOT ^38^ and further refined in PHENIX ^39^. The final refinement statistics are summarized in Table S1. Structural figures were prepared by PyMOL. Epitope and paratope residues, as well as their interactions, were identified by PISA (http://www.ebi.ac.uk/pdbe/prot_int/pistart.html) at the European Bioinformatics Institute.

### Pharmacokinetics (PK) of Nb22-Fc *in vivo*

Purified Nb22-Fc were injected via intranasal (*i.n*.) into BALB/c mice (Qing Long Shan Animal Center, Nanjing, China) at a dose of 10 mg/kg. The concentration of Nb22-Fc in serum was measured by ELISA. The T1/2 of Nb22-Fc was caculated as ln (2)/k, where k is a rate constant expressed reciprocally of the x axis time units by the plateau followed one phase decay or one phase decay equation in the GraphPad software.

### Spatial distribution of Nb22-Fc *in vivo*

Spatial distribution of Nb22-Fc were conducted following our previously published protocol^25^, with minor modifications. Nb22-Fc labeled with far infrared dye YF^®^750 SE (US EVERBRIGHT INC, YS0056) were named as Nb22-YF750. 10 mg/kg Nbs-YF750 were administered via intranasal into nude mice (18-22g, Qing Long Shan Animal Center, Nanjing, China). Images were recorded at Ex:740 nm/Em:780 nm by NightOWL LB 983 (Berthold, Germany) at the indicated time point. Images were analyzed using Indigo imaging software Ver. A 01.19.01.

### Evaluating the efficacy of Nb22-Fc in SARS-CoV-2 infected hACE2 mice

The efficacy of Nb22-Fc against SARS-CoV-2 were evaluated according to our previously published protocol^25^, with minor modifications. In brief, A total of 43 8-week-old male transgenic hACE2 mice (C57BL/6J) (cat.# T037630, GemPharmatech Co., Ltd., Nanjing, China) were challenged with 1× 10^5^ PFU SARS-CoV-2 Delta variant (CRST: 1633.06.IVCAS 6.7593) per mouse. The mice were randomly divided into nine groups (n=3-6) for either prophylactic or therapeutic evaluation, as described in Figure 5A. Mice without any treatment and challenge were taken as blank control (No SARS-CoV-2, n=3). Mice challenged with SARS-CoV-2 were taken as infection control (SARS-CoV-2, n=5). 250 μg/mouse (average of 10 mg/kg) SNB02 (Y-Clone, China), an anti-SFTSV antibody constructed by Nb fused with human Fc1 (Nb-Fc) ^32^, was intranasally injected 1 hour (h) after infection and was taken as an isotype antibody treated control (Control Nb, n=3). For the prophylactic group, mice were intranasally injected with Nb22-Fc at a dose of 250 μg/mouse (average of 10 mg/kg) at 7 days (d), 5d, 3d, 1d before infection (named as −7d Nb22, −5d Nb22, −3d Nb22, −1d Nb22, respectively, n=5-6). For the therapeutic group, mice were intranasally injected with Nb22-Fc at a dose of 250 μg/mouse (average of 10 mg/kg) 1 h or 24 h after infection (named as 1h Nb22 and 1d Nb22, n=5, respectively). Body weight of mouse was recorded daily. Given that hACE2 transgenic mice typically clear virus within 5-7 days after SARS-CoV-2 infection^40^, the mice were sacrificed at 4 days post infection (dpi). Subsequently, lung tissues were harvested for viral load determination and tissue sections for immunofluorescence (IF) and hematoxylin and eosin (H&E) staining. All experiments were conducted in a Biosafety Level 3 (BSL-3) facility.

### Viral load measurement by quantitative RT-PCR

Viral load was measured by quantitative real-time PCR (qRT-PCR) on RNA extracted from the supernatant of lung homogenates as reported previously ^41^. Briefly, lung homogenates were prepared by homogenizing perfused lung using an electric homogenizer. The inactivated samples were transferred from the BSL-3 to BSL-2 laboratory and total RNA was extracted from the collected supernatant. Each RNA sample was reverse transcribed to 50 μl cDNA with HiScript II Q RT SuperMix for qPCR (+gDNA wiper) (R223-01). 5 μl cDNA was added into a 25 μl qRT-PCR reaction containing the ChamQ SYBR qPCR Master Mix (High ROX Premixed) (Q341-02, Vazyme Biotech, China) and primers designed to target the nucleocapsid protein of SARS-CoV-2 (5 ′ -GGGGAACTTCTCCTGCTAGAAT −3 ′ and 5 ′ - CAGACATTTTGCTCTCAAGCTG −3′). The samples were run in triplicate on an ABI 7900 Real-Time System (Applied Biosystems, Thermo Fisher Scientific). The following cycling conditions were performed: 1 cycle of 50 °C for 2 min, 1 cycle of 95 °C for 10 min, and 40 cycles of 95 °C for 15 sec and 58 °C for 1 min.

### Immunofluorescence staining of SARS-CoV-2-infected cells and H&E staining in tissues

The lung tissues were immersed in 10% neutral buffered formalin (cat.# Z2902, Sigma) for 24 h. After the formalin fixation, the tissues were placed in 70% ethanol (Merck) and subsequently embedded with paraffin. Tissue sections (5-μm thick) were prepared for H&E staining and immunofluorescence staining for SARS-CoV-2 detection using the Coronavirus nucleocapsid protein (NP) antibody (cat. 40143-MM05, Sino Biological). Images were collected under a Pannoramic MIDI system (3DHISTECH, Thermo) using Pannoramic scanner software and analyzed by ImageJ (NIH).

### Quantification and statistical analysis

All statistical analyses were carried out using GraphPad Prism 8.01 software (GraphPad) or OriginPro 8.5 software (OriginLab). ANOVA or Mann-Whitney test was performed for group comparisons. *P* < 0.05 was considered as statistically significant with mean ±SEM or mean ±SD.

### Study approval

The study and the protocol for this research were approved by the Center for Public Health Research, Medical School, Nanjing University. All animal experimental procedures without infection were approved by the Committee on the Use of Live Animals by the Ethics Committee of Nanjing University. All animals infected by SARS-CoV-2 were performed in Biosafety Level 3 animal facilities in accordance with the recommendations for care and use of the Institutional Review Board of Wuhan Institute of Virology of the Chinese Academy of Sciences (Ethics Number: WIVA11202111). All the authors declare their compliance with publishing ethics.

## Author contributions

XW conducted most experiments, analyzed the data and wrote the draft manuscript. LC conducted all the neutralization experiments. LZ, BH, MJ, SX, HS, DZ, LL, WN provided technical assistance. YW, SM and SY conducted the structural analysis. FN, YL, HH, QH and YL, evaluated the efficacy of Nb22 in SARS-CoV-2 infected transgenic hACE2 mice. ZW designed the study, directed and revised the manuscript. All authors critically reviewed the draft manuscript and approved the final version.

## Acknowledgements

We thank Prof. Guo. for providing the plasmid of RBD. The X-ray data were collected using Shanghai Synchrotron Radiation Facility on BL18U1 and BL02U1. The efficacy of Nb22 in SARS-CoV-2 infected transgenic hACE2 mice was evaluated in Wuhan National Biosafety Laboratory, Chinese Academy of Sciences. This work was supported by National Science Foundation of China (NSFC) (No. 81803414 to X.W., 31970149 to Z.W.), the Major Research and Development Project (2018ZX10301406 to Z.W.), Ministry of Science and Technology (2020YFA0908500 to S.Y.), the National Natural Science Foundation of China (31971127 to S.Y. and 81801998 to Y.W.), Tianjin Natural Science Foundation (20JCQNJC01570 to Y.W.), Nanjing University-Ningxia University Collaborative Project (Grant# 2017BN04 to Z.W.), Jiangsu Province Natural Science Foundation for Young Scholar (Grant# BK20170653 to X.W.), Key Natural Science Foundation of Jiangsu Province (Grant# ZDA2020014 to X.W.), the Fundamental Research Funds for the Central Universities Grant# 0214-14380523 to X.W.) Jiangsu province “Innovative and Entrepreneurial talent” and Six Talent Peaks Project of Jiangsu Province, the Emergency Prevention and Control Capacity Program for New Severe Infectious diseases of National Institute for Viral Disease Control and Prevention, and the 135 Strategic Program of Chinese Academy of Sciences, the Science and Technology Innovation Committee of Shenzhen Municipality (JCYJ20180228162229889 to L.C.), and the National Science Foundation of China (No. 31970172 to Y.L.).

## Supplemental Materials

**Supplemental Figure 1. Nb22 blocks the binding of hACE2 to WH01 RBD.** (**A**) Overlap of Nb22 and hACE2 binding sites on WH01 RBD. hACE2 binding site on WH01 RBD is shown in cyan line. Nb22 binding site is shown in pink line. The overlap region is represented by ellipses with dashed lines. (**B**) The loop (V102-Y117) of Nb22 is clashed with the two helixes on N-terminal of hACE2. The loop is colored in red and helixes are colored in green.

**Supplemental Figure 2. Spational distribution of Nb22 labeled with dye YF^®^750 SE.** Mice were dissected and detected by NightOwl LB 983 after 200 μg Nb22-YF^®^750 SE infusion into mice as indicated in figure 5C. The fluorescence intensity was measured at 2 hours (0d-Nb22), 7d (7d-Nb22), 14d (14d-Nb22), 21d 21d-Nb22 and 28d (8d-Nb22) after infusion of Nb22 via i.n., respectively. Blank, blank mice without infusion of any antibody, was taken as blank control. The fluorescence intensity of various organs including trachea (Tr), lung (Lu), heart (H), stomach(St), intestine (In), liver(Li), spleen (Sp), kidney (Ki), bladder (B), were analyzed

**Supplemental Figure 3. Body weight of mice.** Body weight of mice in figure 6 was recorded at the indicated time point.

**Supplemental Table 1**. Data collection and refinement statistics

**Supplemental Table 2**. Residues contributed to interaction between Nb22 and RBD were identified by PISA at the European Bioinformatics Institute.

