## Supplemental information for "Short-term instantaneous prophylaxis and efficient treatment against SARS-CoV-2 in hACE2 mice conferred by an intranasal nanobody (Nb22)"

1. Center for Public Health Research, Medical School, Nanjing University, Nanjing, P.R. China.
2. Department of Antibody, Abrev Biotechnology Co., Ltd. Nanjing, P.R. China.
3. Tianjin Key Laboratory of Function and Application of Biological Macromolecular Structures, School of Life Sciences, Tianjin University, Tianjin, P.R. China.
4. Institute for Hepatology, National Clinical Research Center for Infectious Disease, Shenzhen Third People's Hospital, Shenzhen, P.R. China.
5. State Key Laboratory of Virology, Wuhan Institute of Virology, Center for Biosafety Mega-Science, Chinese Academy of Sciences, Wuhan, P.R.China
6. Savaid Medical School, University of Chinese Academy of Sciences, Beijing, P.R. China
7. School of Life Sciences, Ningxia University, Yinchuan, P.R. China.
8. Department of Antibody, Y-clone Medical Science Co. Ltd. Suzhou, P.R. China.
9. Institute for Infection and Immunity, St George's University of London, London, UK
10. Life Sciences Institute, Zhejiang University, Zhejiang, P.R. China.
11. Jiangsu Key Laboratory of Molecular Medicine, Medical School, Nanjing University, Nanjing, P.R. China.
12. State Key Laboratory of Analytical Chemistry for Life Science, Nanjing University, Nanjing, P.R. China.
13. Lead contact

†These authors contributed equally to this work.

Mailing address: 22 Hankou Road, Nanjing, Jiangsu 210093. China

**Declaration of interests:** The authors have declared that no conflict of interest exists.

**This file includes:**

**Supplemental Figure 1. Nb22 blocks the binding of hACE2 to WH01 RBD.** (**A**) Overlap of Nb22 and hACE2 binding sites on WH01 RBD. hACE2 binding site on WH01 RBD is shown in cyan line. Nb22 binding site is shown in pink line. The overlap region is represented by ellipses with dashed lines. (**B**) The loop (V102-Y117) of Nb22 is clashed with the two helixes on N-terminal of hACE2. The loop is colored in red and helixes are colored in green.

**Supplemental Figure 2. Spational distribution of Nb22 labeled with dye YF®750 SE.** Mice were dissected and detected by NightOwl LB 983 after 200 μg Nb22-YF®750 SE infusion into mice as indicated in figure 5C. The fluorescence intensity was measured at 2 hours (0d-Nb22), 7d (7d-Nb22), 14d (14d-Nb22), 21d（21d-Nb22）and 28d（8d-Nb22） after infusion of Nb22 via i.n., respectively. Blank, blank mice without infusion of any antibody, was taken as blank control. The fluorescence intensity of various organs including trachea (Tr), lung (Lu), heart (H) , stomach(St), intestine (In), liver(Li), spleen (Sp), kidney (Ki), bladder (B), were analyzed.

**Supplemental Figure 3. Body weight of mice.** Body weight of mice in figure 6 was recorded at the indicated time point.

**Supplemental Table 1**. Data collection and refinement statistics

**Supplemental Table 2**. Residues contributed to interaction between Nb22 and RBD were identified by PISA at the European Bioinformatics Institute.

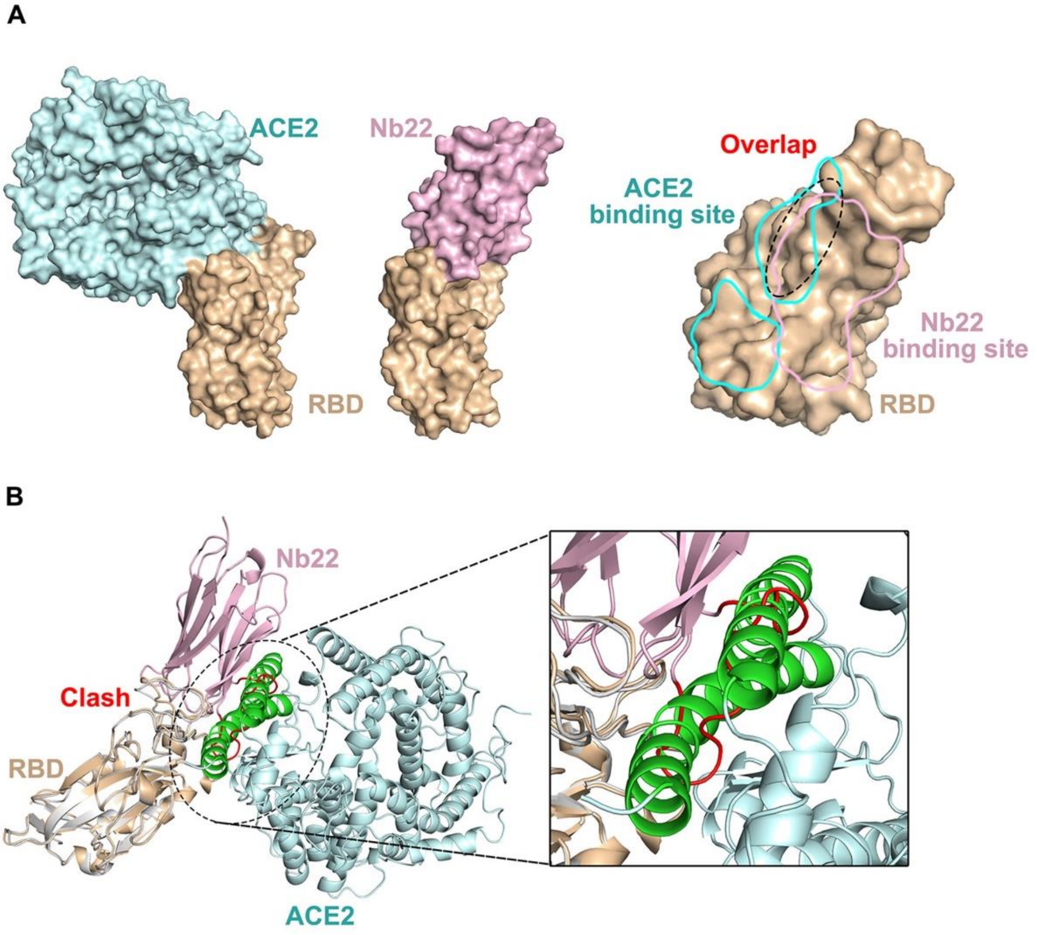


**Supplemental Figure 1. Nb22 blocks the binding of hACE2 to WH01 RBD.** (**A**) Overlap of Nb22 and hACE2 binding sites on WH01 RBD. hACE2 binding site on WH01 RBD is shown in cyan line. Nb22 binding site is shown in pink line. The overlap region is represented by ellipses with dashed lines. (**B**) The loop (V102-Y117) of Nb22 is clashed with the two helixes on N-terminal of hACE2. The loop is colored in red and helixes are colored in green.

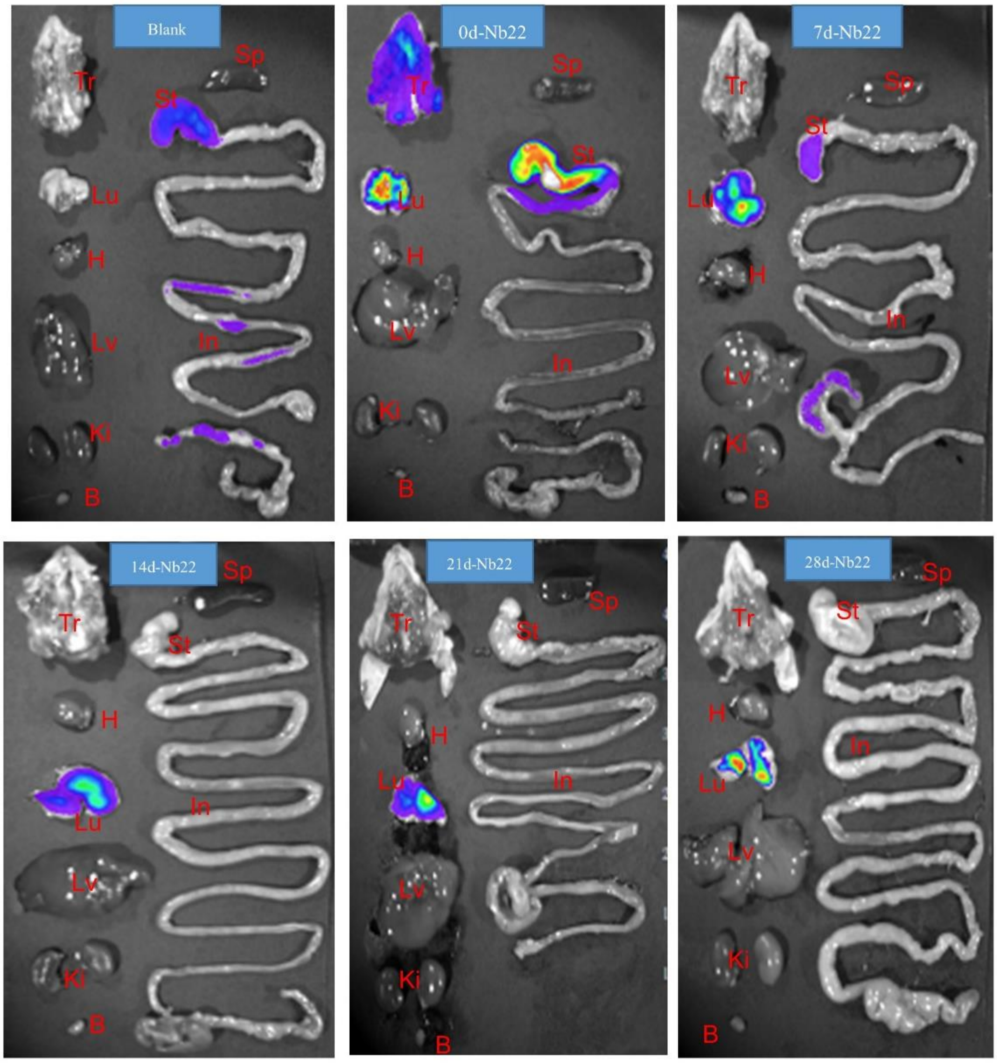


**Supplemental Figure 2. Spational distribution of Nb22 labeled with dye YF®750 SE.** Mice were dissected and detected by NightOwl LB 983 after 200 μg Nb22-YF®750 SE infusion into mice as indicated in figure 5C. The fluorescence intensity was measured at 2 hours (0d-Nb22), 7d (7d-Nb22), 14d (14d-Nb22), 21d（21d-Nb22）and 28d（8d-Nb22） after infusion of Nb22 via i.n., respectively. Blank, blank mice without infusion of any antibody, was taken as blank control. The fluorescence intensity of various organs including trachea (Tr), lung (Lu), heart (H) , stomach(St), intestine (In), liver(Li), spleen (Sp), kidney (Ki), bladder (B), were analyzed.

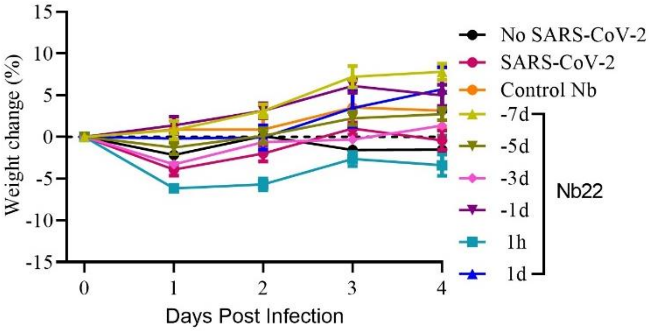


**Supplemental Figure 3. Body weight of mice.** Body weight of mice in figure 6 was recorded at the indicated time point.

**Supplemental Table 1. Data collection and refinement statistics**

| Parameters | WH01 RBD-Nb22 | Delta RBD-Nb22 |
|---|---|---|
| X-ray Source | BL18U1 | BL02U1 |
| Wavelength (Å) | 0.97915 | 0.97918 |
| Space group | $P2_12_12_1$ | $C2$ |
| Unit cell parameters (Å) | $a$=73.8, $b$=88.7, $c$=172.5 | $a$=151.8, $b$=108.5, $c$=115.3 <br> $\alpha$=90.0, $\beta$=126.2, $\gamma$=90.0 |
| Resolution range (Å) | 50.0-2.7 (2.75-2.7)* | 93.0-2.9 (3.0-2.9)* |
| Unique reflections | 32,672 (1,579) | 30,823 (3,106) |
| Completeness (%) | 99.8 (99.2) | 93.8 (95.7) |
| Redundancy | 7.2 (4.8) | 3.2 (3.1) |
| I/σ(I) | 12.3 (1.7) | 6.9 (2.5) |
| $R_{merge}$ (%) | 13.7 (64.4) | 11.0 (41.6) |
| $R_{meas}$ (%) | 14.8 (72.2) | 13.1 (50.0) |
| $R_{pim}$ (%) | 5.5 (32.1) | 7.1 (27.4) |
| $CC_{1/2}$ | 0.999 (0.763) | 0.985 (0.836) |
| **Refinement statistics** | | |
| Resolution range (Å) | 37.23- 2.7 (2.796-2.7) | 41.2-2.9 (3.0-2.9) |
| Reflections used in refinement | 31,683 (3,093) | 30,797 (3,104) |
| Reflections used for R-free | 1,663 (162) | 1,495 (140) |
| $R_{work}$ (%) | 19.6 (27.3) | 20.5 (27.2) |
| $R_{free}$ (%) | 24.4 (35.4) | 25.5 (32.1) |
| Number of non-hydrogen atoms | 7,710 | 7,539 |
| Protein | 972 | 972 |
| Solvent | 155 | / |
| Ligand | 13 | / |
| Average B-factors | 59.1 | 85.7 |
| Protein | 59.4 | 85.7 |
| Solvent | 40.7 | / |
| Ligand | 52.0 | / |
| r.m.s. deviations | | |
| Bond lengths (Å) | 0.003 | 0.009 |
| Bond angles (°) | 0.54 | 1.11 |
| Ramachandran | | |
| Favored (%) | 94.9 | 94.3 |
| Allowed (%) | 5.1 | 5.7 |
| Outliers (%) | 0.0 | 0.0 |

*Numbers in the brackets are for the highest resolution shell.

**Supplemental Table 2. Residues contributed to interaction between Nb22 and RBD were**

**identified by PISA at the European Bioinformatics Institute.**

| Nb22 | Distance (Å) | WH01 RBD | Distance (Å) | Delta RBD |
|---|---|---|---|---|
| GLY 1[O] | 2.9 | ASN 450[ND2] | 2.8 | ASN 450[ND2] |
| THR 30[N] | 2.9 | SER 494[OG] | 3.0 | SER 494[OG] |
| THR 30[OG1] | / | ARG 452[NE] | 2.9 | ARG 452[NE] |
| SER 33[OG] | / | GLN 493[OE1] | 2.8 | GLN 493[OE1] |
| SER 33[OG] | 2.6 | SER 494[N] | 3.0 | SER 494[N] |
| ASN 57[ND2] | 2.9 | GLY 485[O] | 2.9 | GLY 485[O] |
| SER 75[N] | 2.4 | GLU 484[OE2] | 2.6 | GLU 484[OE2] |
| SER 75[OG] | 2.8 | GLU 484[OE1] | 2.8 | GLU 484[OE1] |
| TYR 119[OH] | 2.8 | GLY 446[O] | 2.8 | GLY 446[O] |
